# *In silico* analyses reveal new putative Breast Cancer RNA-binding proteins

**DOI:** 10.1101/2020.01.08.898965

**Authors:** Santiago Guerrero, Andrés López-Cortés, Jennyfer M. García-Cárdenas, Isaac Armendáriz-Castillo, Ana Karina Zambrano, Alberto Indacochea, Andy Pérez-Villa, Verónica Yumiceba, Patricia Guevara-Ramírez, Andrea-Jácome-Alvarado, Paola E. Leone, César Paz-y-Miño

## Abstract

Breast cancer (BC) is the leading cause of cancer-associated death among women worldwide. Despite treatment efforts, advanced BC with distant organ metastases is considered incurable. A better understanding of BC molecular processes is therefore of great interest to identify new therapeutic targets. Although large-scale efforts, such as The Cancer Genome Atlas (TCGA), have completely redefined cancer drug development, diagnosis, and treatment, additional key aspects of tumor biology remain to be discovered. In that respect, post-transcriptional regulation of tumorigenesis represents an understudied aspect of cancer research. As key regulators of this process, RNA-binding proteins (RBPs) are emerging as critical modulators of tumorigenesis but only few have defined roles in BC. To unravel new putative BC RBPs, we have performed *in silico* analyses of all human RBPs in three major cancer databases (TCGA-Breast Invasive Carcinoma, the Human Protein Atlas, and the Cancer Dependency Map project) along with complementary bioinformatics resources (STRING protein-protein interactions and the Network of Cancer Genes 6.0). Thus, we have identified six putative BC progressors (MRPL13, SCAMP3, CDC5L, DARS2, PUF60, and PLEC), and five BC suppressors RBPs (SUPT6H, MEX3C, UPF1, CNOT1, and TNKS1BP1). These proteins have never been studied in BC but show similar cancer-associated features than well-known BC proteins. Further research should focus on the mechanisms by which these proteins promote or suppress breast tumorigenesis, holding the promise of new therapeutic pathways along with novel drug development strategies.

## Introduction

Breast cancer (BC) is the leading cause of cancer-associated death (15%: 626,679 cases) and the most commonly diagnosed cancer (24%: 2,088,849 cases) among women worldwide^1^. BC is characterized by a complex interaction between environmental factors and biological aspects, such as gene deregulation, hormone disruption or ethnicity^2–4^. Despite subtype-specific treatment efforts, advanced BC with distant organ metastases is considered incurable^2^. Therefore, a better understanding of BC molecular processes is still of great interest to identify new therapeutic targets.

Current oncological research generates large-scale datasets that harbor essential aspects of tumor biology. For instance, The Cancer Genome Atlas (TCGA), with over 2.5 petabytes of data, has molecularly characterized over 20,000 patient samples covering 33 cancer types^5–10^. Also, The Cancer Dependency Map (DepMap) project, using loss-of-function genetic screens, has identified essential genes for cancer proliferation and survival *ex vivo*^11–13^. Additionally, The Human Protein Atlas (HPA) constitutes a comprehensive resource to explore the human proteome in healthy and tumor tissues^14–16^. Although these datasets have completely redefined cancer drug development, diagnosis, and treatment, additional key aspects of tumor biology remain to be discovered. In that respect, post-transcriptional regulation of tumorigenesis represents an understudied aspect of cancer research^17^.

As key regulators of post-transcription, RNA binding proteins (RBPs) are particularly relevant due to their implication in every post-transcriptional step of gene expression: RNA splicing, transport, stability, translation, and localization. As a result, genomic alterations of these proteins lead to dysfunctional cellular processes; indeed, RBPs are emerging as critical regulators of tumorigenesis but only few have well-defined roles in BC^18–26^. To date, 1,393 RBPs have been experimentally identified from human RNA interactomes^27^. Despite efforts to understand their role in cancer^28,29^, an integrated analysis of the aforementioned databases along with other *in silico* approaches is still missing in BC. To shed light on this matter, we analyzed and integrated RBPs genomic alterations, protein–protein interaction (PPI) networks, immunohistochemical profiles, and loss-of-function experiments to unravel new putative BC RBPs.

## Results

### An Overview of RBPs genomic alterations in BC

To globally assess the potential role of RBPs in BC versus well-known BC genes, we first interrogated the Breast Invasive Carcinoma (TCGA, PanCancer Atlas)^5–10^ database for genomic alterations of RBPs (n=1392), BC genes (n=171)^30^ and non-cancer genes (n=170)^31^ (Supp. Table 1). As shown in Fig. 1A, both genomic alterations frequencies of RBPs and BC genes were significantly higher than the one observed for non-cancer genes. Interestingly, RBPs present a similar amount of genomic alterations than BC genes (Fig. 1A), highlighting the putative role of RPBs in BC.

**Figure 1.**
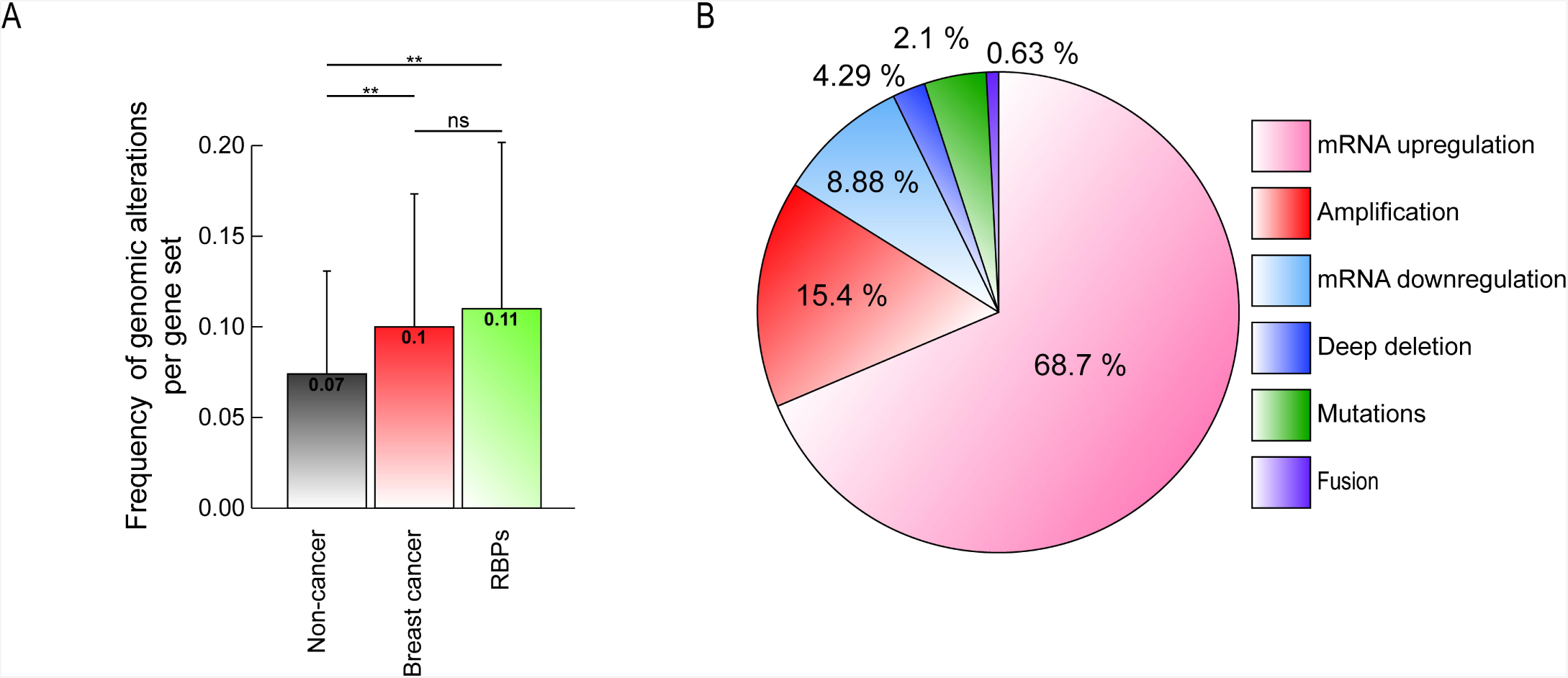
Genomic alterations of RNA-binding proteins (RBPs) in BC. (A) Frequency of genomic alterations per gene set (non-cancer genes [n=170], BC genes [n=171], and RBPs [n=1392]) using the Breast Invasive Carcinoma (TCGA, PanCancer Atlas)^5–10^ database. Genomic alterations per patient were corrected by the number of genes; a Mann–Whitney U test was used to compare genomic alterations between gene sets. ** = highly significant; ns = not significant. (B) A pie chart describing RBPs genomic alterations types.

To obtain insights into how these proteins are altered in BC, we catalogued their genomic alteration types. As shown in Fig. 1B and Supp. Table 2, most genomic alterations are related to an overrepresentation of the mRNA (68.7%) or gene loci (15.4%).

### Identification of highly altered BC RBPs

To identify BC-related RBPs, we next interrogated The Network of Cancer Genes 6.0 (NCG6)^30^ for RBPs having known or predicted cancer driver roles. NCG6 harbors the most recent catalog of cancer driver genes^30^. Thus, we identified 225 RBPs: 14 implicated in BC (2 oncogenes, 4 tumor suppressors, and 8 unknown), indicating that these proteins remain poorly studied in breast carcinogenesis, and 211 related to other cancer types (21 oncogenes, 24 tumor suppressors, and 166 unknown) (Fig. 2A, Supp. Table 3).

**Figure 2.**
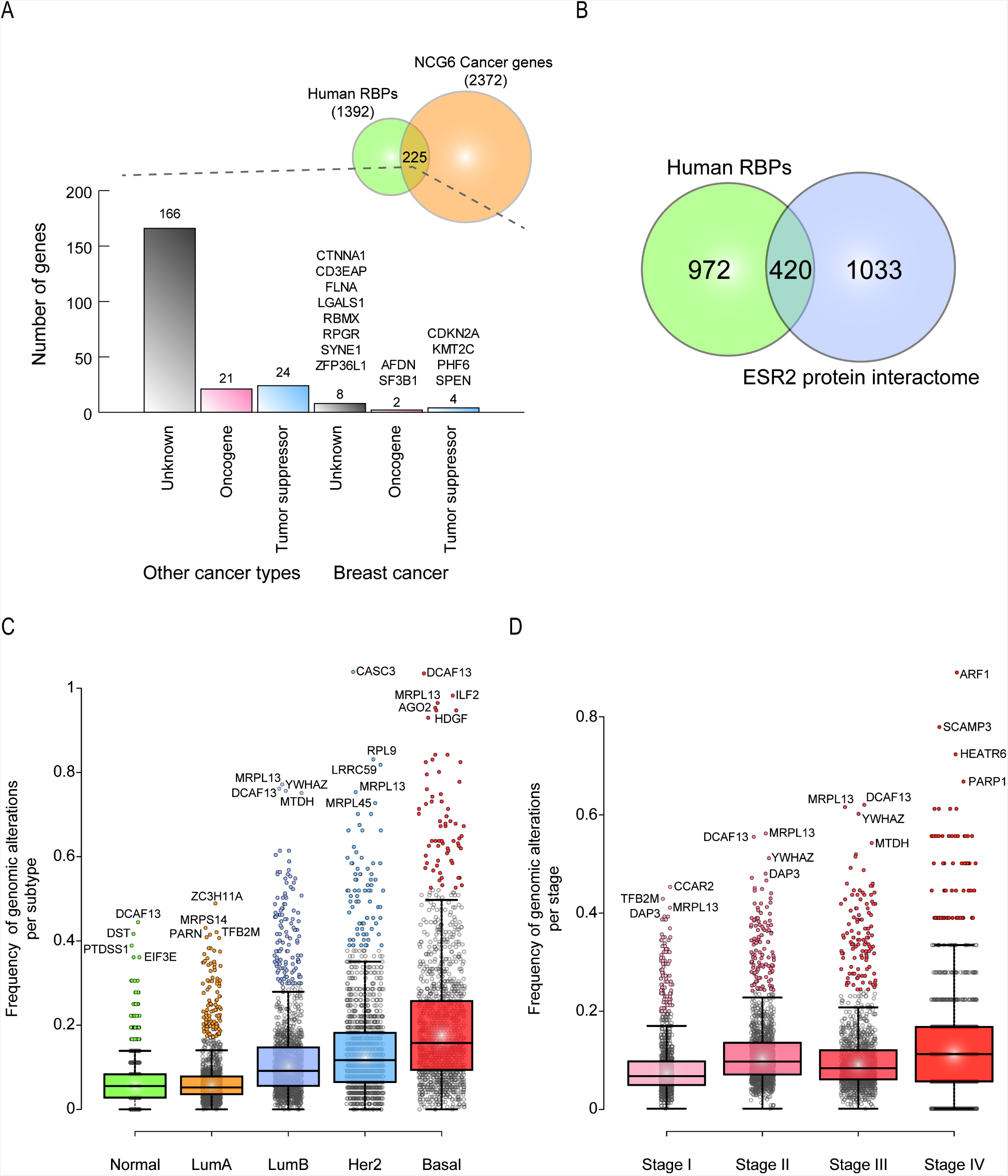
Identification of Highly Altered BC RBPs. (A) An histogram describing the status of RBPs in the Network of Cancer Genes 6.0 (NCG6). (B) A Venn diagram depicting the relationship between RBPs and the estrogen receptor 2 (ESR2) protein interactome. RBPs genomic alterations per subtype (C) and stage (D) using the Breast Invasive Carcinoma (TCGA, PanCancer Atlas)^5–10^ database are also displayed.

To unravel putative RBPs implicated in tumor progression or suppression, we analyzed RBPs genomic alterations based on their progressor or suppressor profiles. Tumor progressors tend to be overexpressed (mRNA upregulation or genomic amplification), while suppressors are downregulated (mRNA downregulation or genomic deletion)^32^. Gene mutations or fusions have been observed in both scenarios. On this basis, we identified highly altered BC RBPs (Table 1, Supp. Table 2). For instance, MRPL13 molecular and cellular functions have never been studied in BC; however, MRPL13 has shown to interact with the estrogen receptor 2 (ESR2), a tumor suppressor protein in breast and other cancers^33^. Interestingly, 30% of all human RBPs interact with the tumor suppressor ESR2^33^ (Fig. 2B). We also found known BC progressors and suppressors proteins, such as DAP3^18^, MTDH^19^ or CCAR2^20^, validating our strategy (Table 1, Supp. Table 2). Also, these analyses reveal proteins that have not been related to tumorigenesis, and yet they are highly altered in BC (e.g. TFB2M, C1ORF131 or DDX19A) (Table 1, Supp. Table 2).

**Table 1.**
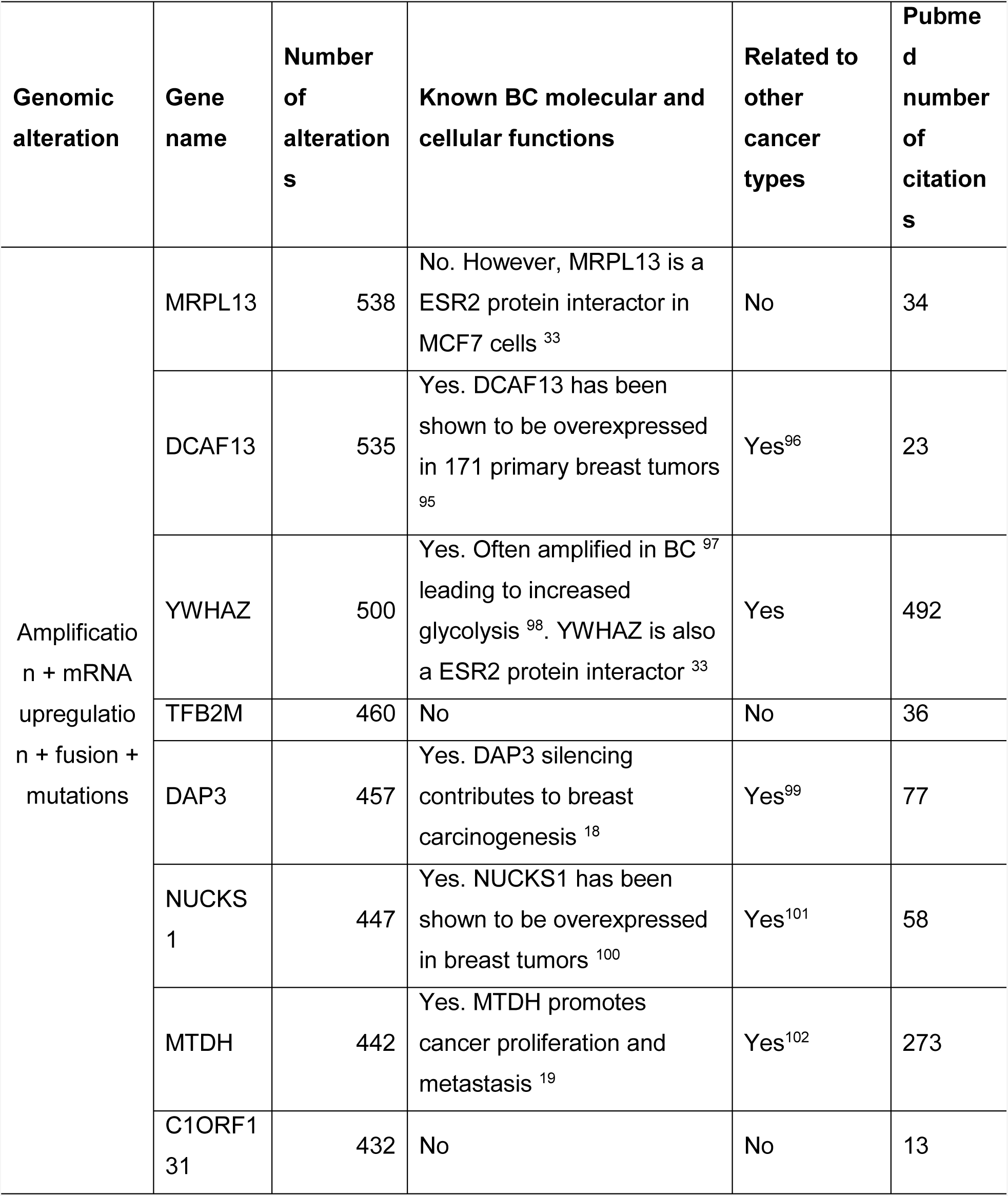
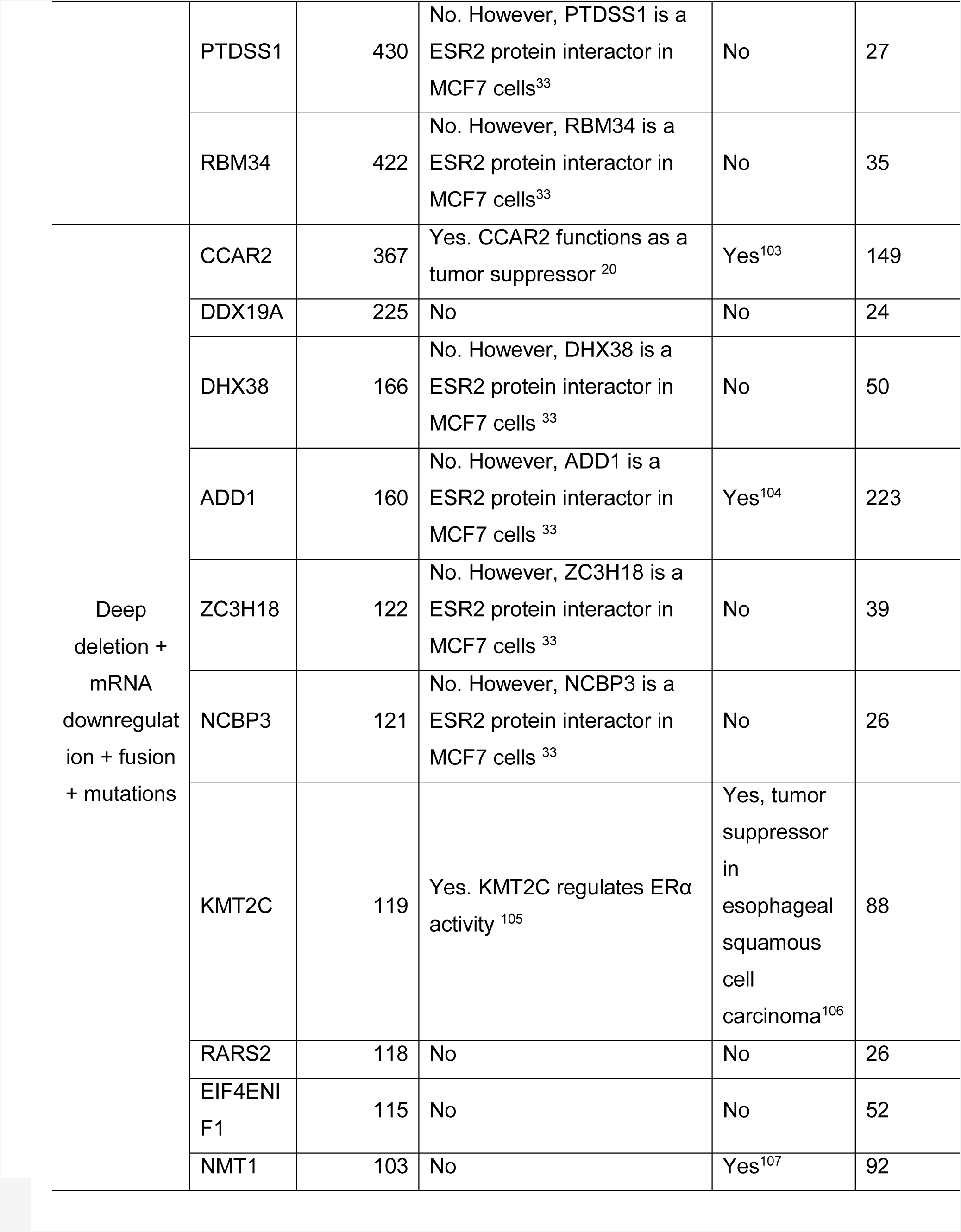
Top ten most altered RBPs in Breast Invasive Carcinoma (TCGA, PanCancer Atlas)

| **Genomic alteration** | **Gene name** | **Number of alterations** | **Known BC molecular and cellular functions** | **Related to other cancer types** | **Pubmed number of citations** |
| --- | --- | --- | --- | --- | --- |
| Amplification + mRNA upregulation + fusion + mutations | MRPL13 | 538 | No. However, MRPL13 is a ESR2 protein interactor in MCF7 cells <sup>33</sup> | No | 34 |
|  | DCAF13 | 535 | Yes. DCAF13 has been shown to be overexpressed in 171 primary breast tumors <sup>95</sup> | Yes<sup>96</sup> | 23 |
|  | YWHAZ | 500 | Yes. Often amplified in BC <sup>97</sup> leading to increased glycolysis <sup>98</sup>. YWHAZ is also a ESR2 protein interactor <sup>33</sup> | Yes | 492 |
|  | TFB2M | 460 | No | No | 36 |
|  | DAP3 | 457 | Yes. DAP3 silencing contributes to breast carcinogenesis <sup>18</sup> | Yes<sup>99</sup> | 77 |
|  | NUCKS1 | 447 | Yes. NUCKS1 has been shown to be overexpressed in breast tumors <sup>100</sup> | Yes<sup>101</sup> | 58 |
|  | MTDH | 442 | Yes. MTDH promotes cancer proliferation and metastasis <sup>19</sup> | Yes<sup>102</sup> | 273 |
|  | C1ORF131 | 432 | No | No | 13 |
|  | PTDSS1 | 430 | No. However, PTDSS1 is a ESR2 protein interactor in MCF7 cells<sup>33</sup> | No | 27 |
|  | RBM34 | 422 | No. However, RBM34 is a ESR2 protein interactor in MCF7 cells<sup>33</sup> | No | 35 |
| Deep deletion + mRNA downregulation + fusion + mutations | CCAR2 | 367 | Yes. CCAR2 functions as a tumor suppressor<sup>20</sup> | Yes<sup>103</sup> | 149 |
|  | DDX19A | 225 | No | No | 24 |
|  | DHX38 | 166 | No. However, DHX38 is a ESR2 protein interactor in MCF7 cells<sup>33</sup> | No | 50 |
|  | ADD1 | 160 | No. However, ADD1 is a ESR2 protein interactor in MCF7 cells<sup>33</sup> | Yes<sup>104</sup> | 223 |
|  | ZC3H18 | 122 | No. However, ZC3H18 is a ESR2 protein interactor in MCF7 cells<sup>33</sup> | No | 39 |
|  | NCBP3 | 121 | No. However, NCBP3 is a ESR2 protein interactor in MCF7 cells<sup>33</sup> | No | 26 |
| | KMT2C | 119 | Yes. KMT2C regulates ERα activity<sup>105</sup> | Yes, tumor suppressor in esophageal squamous cell carcinoma<sup>106</sup> | 88 |
|  | RARS2 | 118 | No | No | 26 |
|  | EIF4ENIF1 | 115 | No | No | 52 |
|  | NMT1 | 103 | No | Yes<sup>107</sup> | 92 |

To further identify key RBPs implicated in BC, we analyzed RBPs genomic alterations by subtype (Normal-like, Luminal A, Luminal B, Her2-enriched, and Basal-like. Supp. Table 4) or staging (stage I to IV; Supp. Table 5). As shown in Fig. 2C, RBPs are highly altered in Basal-like BC compared to other subtypes. Concerning staging, RBPs seem to be more altered in stage IV. Some RBPs reached higher frequencies of genomic alterations per subtype (Fig. 2C, Supp. Table 4) or stage (Fig. 2D, Supp. Table 5). For instance, ARF1, the most altered protein in stage IV (Fig. 2D), has been shown to promote BC metastasis^34^; PARP1 has also been demonstrated to enhance metastasis not only in BC^35^ but also in other cancer types^36^. In contrast, SCAMP3 and HEATR6 presented a similar amount of genomic alterations (Fig. 2D) and none of them has been previously studied in BC.

### RBPs interact with well-known BC proteins

Networking analysis has proven to be useful in identifying RNA regulons and key tumoral proteins^37^. On this basis, we next explored PPIs between RBPs (n=1392) and well-known BC proteins (n=171)^30^ using STRING database^38^. PPIs were obtained from experiments and databases (interaction score=0.9, highest confidence). We identified 113 BC proteins interacting with 398 RBPs (Supp. Table 6). By narrowing down our analysis to experimental interactions only (Fig. 3), we observed two main networks around SF3B1 and CDC5L proteins. According to g:Profiler^39^, proteins interacting with SF3B1 are implicated in RNA splicing (Padj=3.783×10^−34^; GO:0000377), while proteins connected to CDC5L are manly involved in chromatin binding (Padj=1.500×10^−2^; GO:0003682). We also observed proteins with both BC and RNA-binding features present in the two main networks: SF3B1, CTNNA1, RBMX, and SPEN. Additionally, 18 RBPs interact with at least one BC protein. Thus, we identified RBPs that may have a putative role in BC molecular pathways through PPIs.

**Figure 3.**
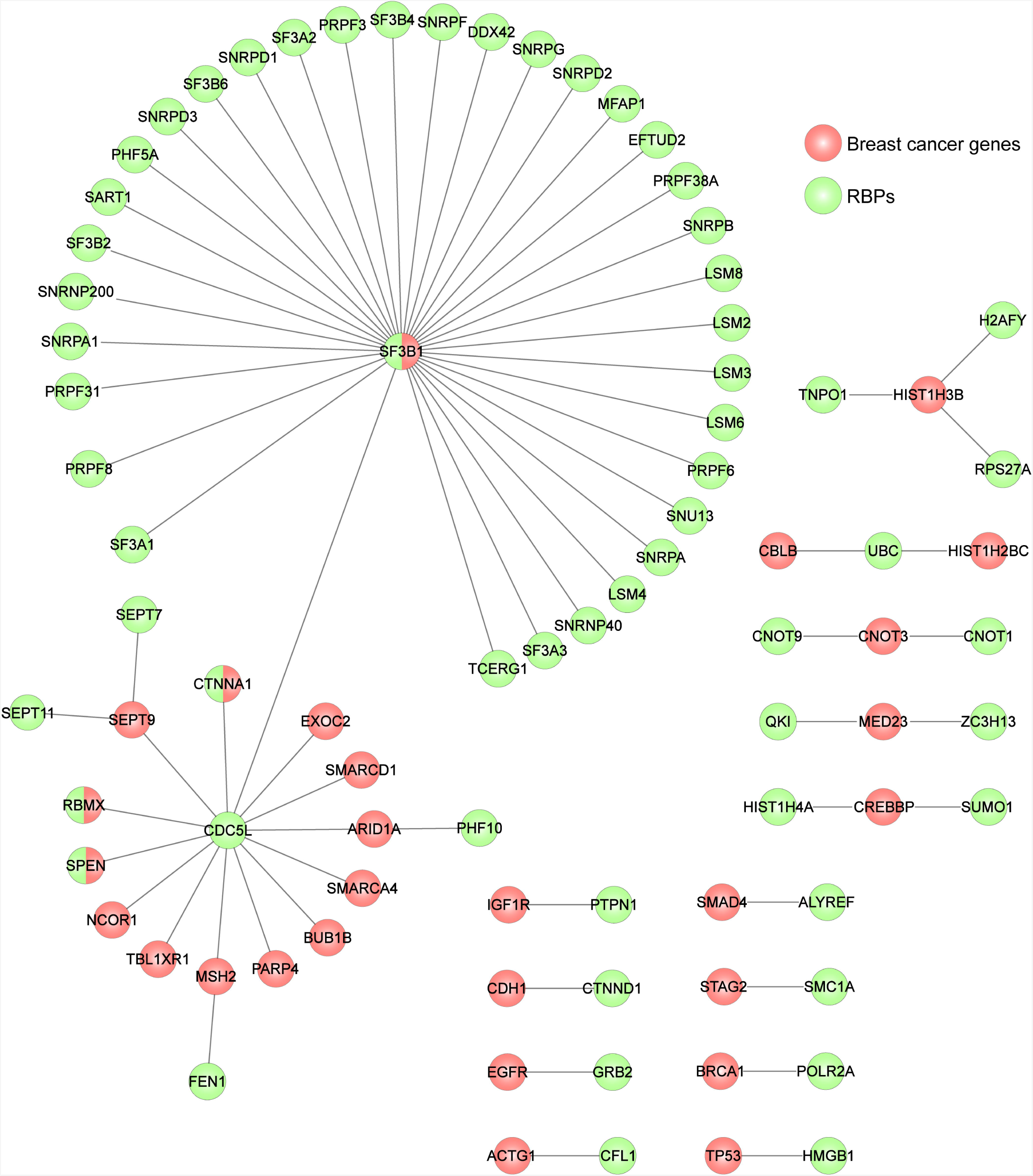
Experimental PPIs between RBPs and well-known BC proteins. An interaction network, constructed using STRING database and Cytoscape 3.7.1 platform, is presented; red, BC proteins; green, RBPs.

### Identification of differentially expressed RBPs in breast tumor tissues

The Human Protein Atlas (HPA) constitutes^14–16^ a major effort to address protein expression in healthy and tumoral human tissues. We therefore identified RBPs having a differentially protein expression profile in breast tumor tissues. To this end, we compared immunohistochemical levels (not detected, low, medium and high) of 1212 available RBPs between normal and cancerous breast tissues (Fig. 4, Supp. Table 6). Most RBPs presented common immunohistochemical levels between both breast tissues: not detected (n=130), low (n=52), medium (n=366), and high (n=172) (Fig. 4A). Moderate protein expression changes, defined by one variation level (e.g. not detected to low or medium to high) were observed in 406 RBPs.

**Figure 4.**
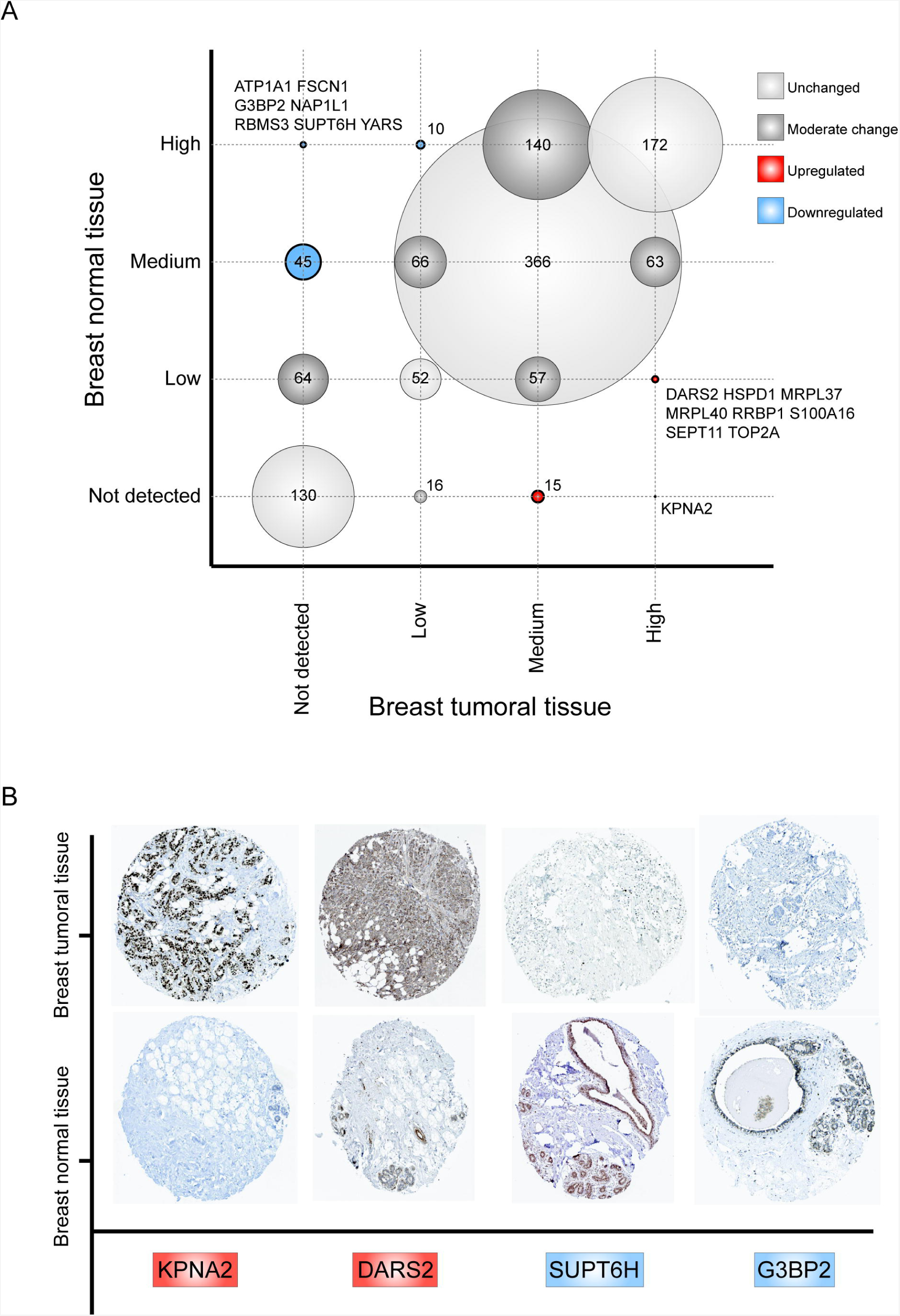
Immunohistochemical protein expression profile of RBPs between healthy and breast tumor tissues. (A) A correlation plot, comparing RBPs immunohistochemical levels between normal and BC tissues, is presented. Circle sizes correlate with the number of RBPs presented in each intersection. (B) Representative immunohistochemical stains of four RBPS (upregulated in red: KPNA2 and DARS2; downregulated in blue: G3BP2 and SUPT6H) on normal and tumor breast tissues according to The Human Protein Atlas (HPA).

To identify RBPs with highly altered protein expression profiles in tumor tissues, we categorized RBPs having a twofold variation level as upregulated or downregulated compared with normal tissues; thus, we identified 24 upregulated and 62 downregulated RBPs (Fig. 4A, Supp. Table 6). As expected, our approach revealed well-known BC proteins, such as KPNA2^21^ or G3BP2^22^, which validate our analysis. KPNA2 is highly expressed in BC tissues (7 out of 12 tumor samples are classified as high) (Fig. 4B, Supp. Table 6). On the contrary, G3BP2 expression is reduced in breast tumor tissues (Fig. 4B, Supp. Table 6). We also observed two RBPs that have never been studied in BC: DARS2 (overexpressed) and SUPT6H (downregulated) (Fig. 4B, Supp. Table 6).

### Exploring RBPs BC dependencies

Most RBPs present numerous genomic alterations (Fig. 1; Fig 2C and D; Supp. Table 2-4), making it difficult to detect essential RBPs for tumor survival. To study RBPs BC dependencies, we analyzed 1288 available RBPs on CERES^11^ and 1290 available RBPs on DEMETER2^12,13^ through the DepMap portal (https://depmap.org/portal/). Both initiatives report loss-of-function screens performed in several human cancer cell lines^11–13^.

Fig. 5A shows the distribution of dependency scores of all available RBPs in 82 (DEMETER2^12,13^) and 28 (CERES^11^) BC cell lines. The genome-scale RNAi loss-of-function screens (DEMETER2^12,13^) identified 90 essential RBPs (Fig. 5A), being SNRPD1, SF3B1, SF3B2, RPL5, ARCN1, EIF3B, RAN, COPB1, RPL14, and VCP (mean dependency scores ranging from −1.3 to −1.5) the top ten essential RBPs for BC survival (Supp. Table 8). On the other hand, genome-scale CRISPR-Cas9 loss-of-function screens (CERES^11^) determined 176 essential RBPs (Fig. 5A), being RAN, HSPE1, SNRNP200, SNRPD1, SARS, EEF2, RPL37, CCT3, KPNB1, and RPL23 (mean dependency scores ranging from −1.5 to −1.8) the top ten essential RBPs for tumor survival (Supp. Table 9). *In toto*, 207 essential RBPs were identified by both computational methods (Fig. 5A; Supp. Table 8-9).

**Figure 5.**
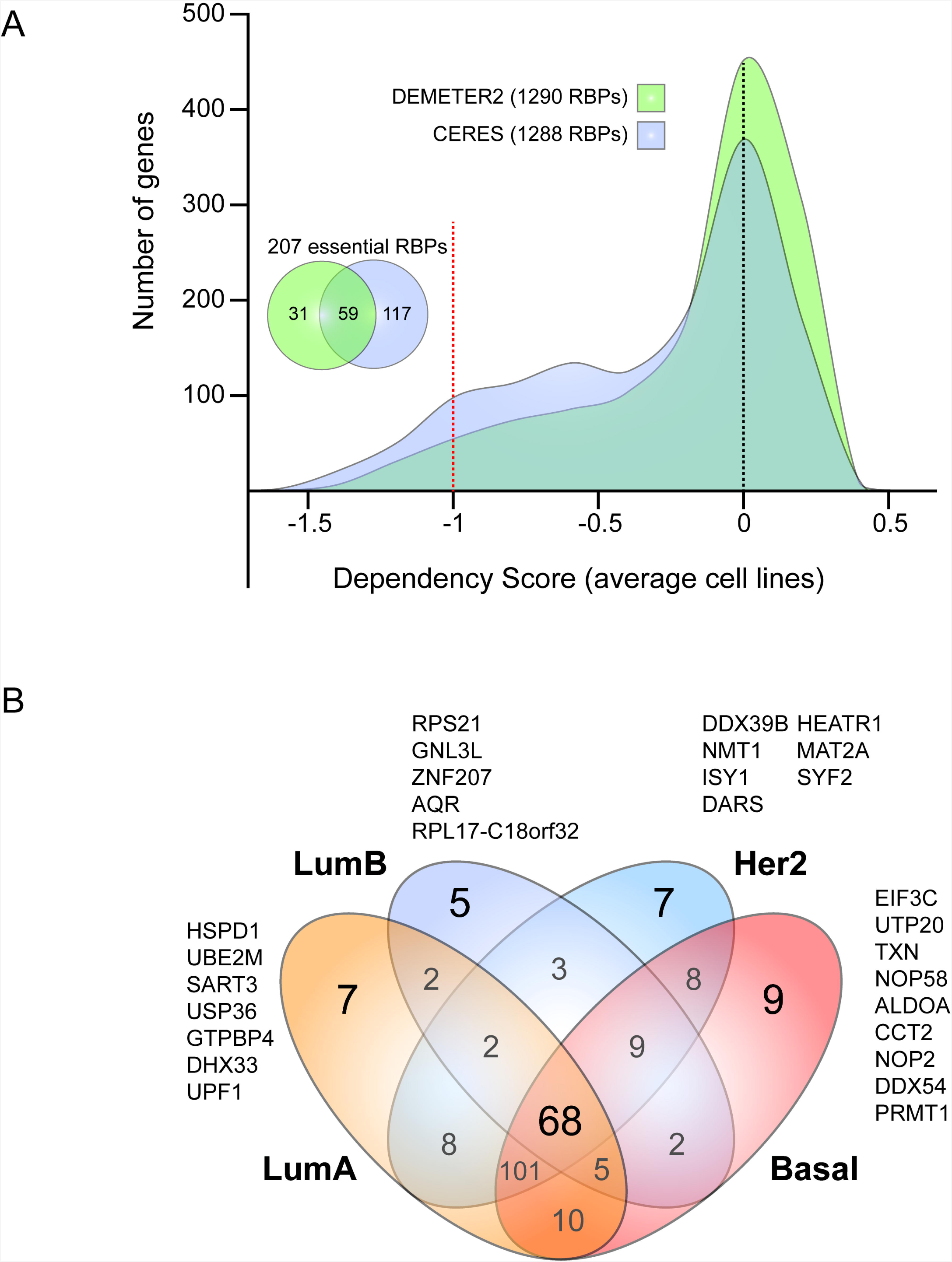
RBPs BC dependencies. (A) The distribution of dependency scores of 1290 (DEMETER2) and 1288 RBPs (CERES) is shown. (B) A Venn diagram comparing BC essential RBPs per subtype is presented.

To identify essential RBPs per BC molecular subtype, we first updated subtypes by merging data from Smith *et al.*^40^, Dai *et al.*^41^, and Kao *et al.*^42^ (Supp. Table 10). Next, we identified and compared 203 LumA, 96 LumB, 206 Her2, and 212 Basal essential RBPs (Fig. 5B; Supp. Table 8-9). Thus, we identified essential RBPs for each BC subtype: 7 LumA (HSPD1, UBE2M, SART3, USP36, GTPBP4, DHX33, and UPF1), 5 LumB (RPS21, GNL3L, ZNF207, AQR, and RPL17-C18orf32), 7 Her2 (DDX39B, NMT1, ISY1, DARS, HEATR1, MAT2A, and SYF2), and 9 Basal (EIF3C, UTP20, TXN, NOP58, ALDOA, CCT2, NOP2, DDX54, and PRMT1) (Fig. 5B).

### Unraveling putative BC RBPs

Cancer-related RBPs control hundreds of tumor mRNAs, interact with well-known cancer driver proteins and appear to be highly altered in cancer genomic databases and tumor tissues^32^. We therefore reasoned that integration of our previous analyses could narrow down the identification of potential BC RBPs.

To this end, we first focused on RBPs having putative tumor progression profiles. Thus, we overlapped our previous results as follows: 1) 348 RBPs belonging to the first quartile of most genomically-altered RBPs concerning tumor progression-related alterations (mRNA upregulation, genomic amplification, gene mutations or fusions); 2) All 398 RBPs presenting PPIs with well-known BC proteins (Supp. Table 6); 3) 160 RBPs having at least one immunohistochemical variation level towards protein overexpression (e.g. not detected to low); And 4) All 207 BC essential RBPs (Fig. 5A; Supp. Table 8-9).

We found five RBPs presenting the aforementioned tumor-associated characteristics: TFRC, KPNB1, PUF60, NSF, and SF3A3 (Fig. 6A). TFRC and KPNB1 have been previously implicated in BC^43–46^, while PUF60 has been associated with colon and non-small cell lung cancer^47,48^. Interestingly, NSF and SF3A3 have never been studied in cancer. We also found 14 RBPs showing high genomic alterations, PPIs with BC proteins, and altered protein expression profiles in tumoral tissues. Although these proteins are not needed for tumor survival *ex vivo* (Supp. Table 8-9), they could be implicated in other tumorigenic processes; indeed, 11 of these RBPs have been described as BC tumor progressors^49–60^. Interestingly, PLEC has not been related with BC, but promotes migration and invasion of neck squamous cell carcinoma^61^. In addition, PRPF3 and MAGOHB have not been linked to cancer before.

**Figure 6.**
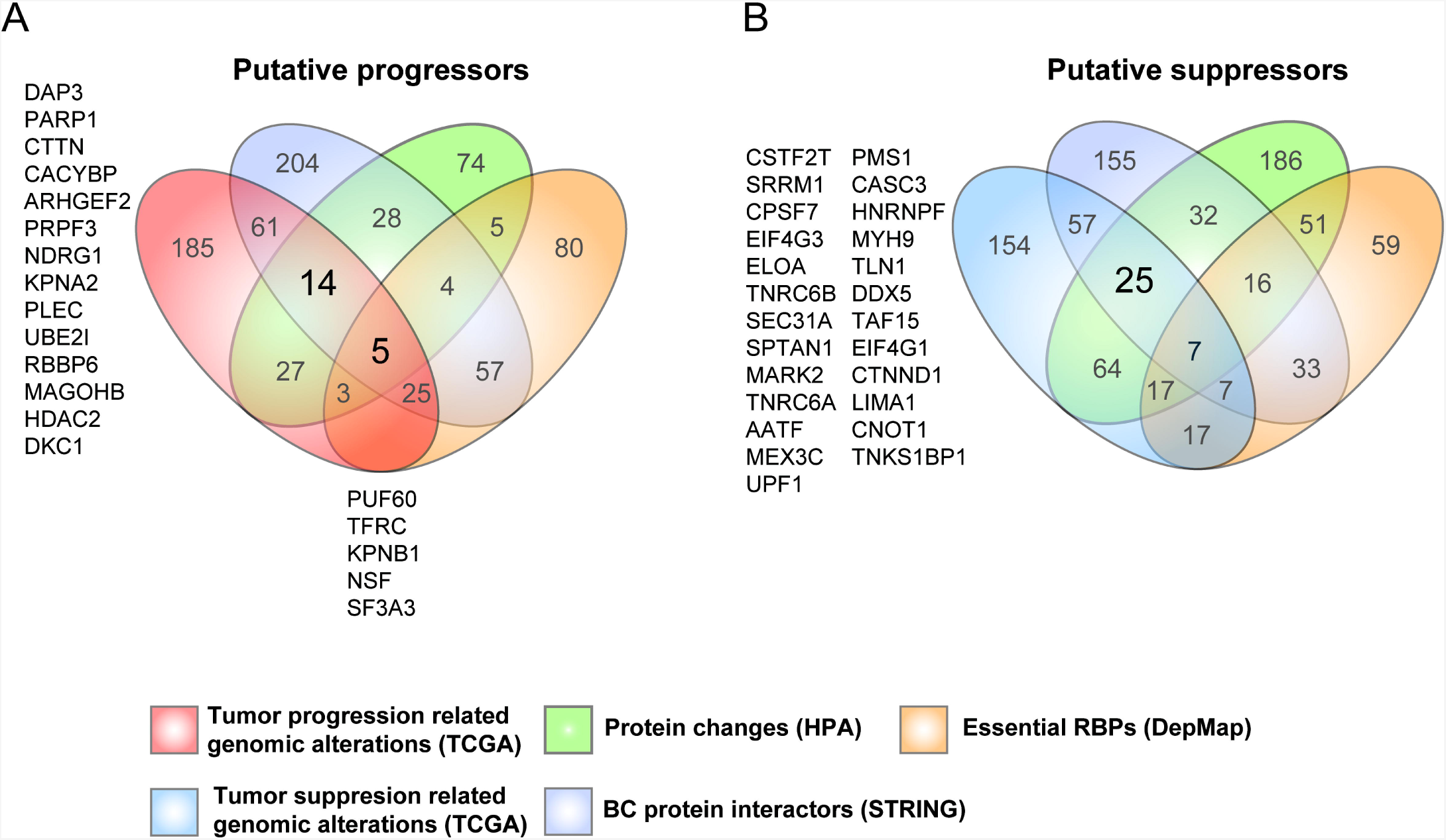
Unraveling putative BC RBPs. (A) A Venn diagram depicting the number of unique and shared RBPs across the four cancer progression profiles. (B) A Venn diagram showing the number of unique and shared RBPs across the four cancer suppression profiles.

Next, we focused on RBPs having putative tumor suppression profiles. In this concern, we compared our afore made analyses as follows: 1) 348 RBPs belonging to the first quartile of most genomically-altered RBPs concerning tumor suppression-related alterations (mRNA downregulation, deep deletions, gene mutations or fusions); 2) All 398 RBPs presenting PPIs with well-known BC proteins (Supp. Table 6); 332 RBPs having at least one immunohistochemical variation level towards protein downregulation (e.g. low to not detected); 4) All 207 BC essential RBPs (Fig. 5A; Supp. Table 8-9). Contrary to our previous analysis, DEMETER2^12,13^ and CERES^11^, which detect loss-of-function proteins, are now used to discriminate between suppressors and progressors RBPs. Thus, we identified 25 RBPs (Fig. 6B): 5 never studied in cancer (CSTF2T, SRRM1, CPSF7, EIF4G3, and ELOA), 6 associated with other cancer types (TNRC6B^62^, SEC31A^63^, SPTAN1^64^, MARK2^65^, TNRC6A^66^, and PMS1^67^), 7 associated with BC (CASC3^68^, HNRNPF^69^, MYH9^70^ TLN1^71^, DDX5^72^, TAF15^73^, and EIF4G1^74^), 3 already described as tumor suppressors in BC (CTNND1^23^, LIMA1^24^, AATF^25^), and 4 identified as tumor suppressors in other cancer types (MEX3C^75^, UPF1^76^, CNOT1^77^, and TNKS1BP1^78^).

## Discussion

Current oncological research generates large-scale datasets that contain undiscovered key aspects of tumor biology, and yet these databases are not fully exploited. *In silico* analyses of these data could therefore lead to the discovery of new cancer proteins^29,79–81^. In this regard, RBPs are emerging as critical regulators of tumorigenesis^17^; however, they remain understudied in cancer research. In the present study, we analyzed and integrated RBPs genomic alterations, PPI networks, immunohistochemical profiles, and loss-of-function experiments to unravel new putative BC proteins.

The results of this research revealed for the first time that RBPs are equally altered than well-known BC proteins (Fig. 1A); this was expected since many RBPs are highly altered across cancer types^29^ and have been *in silico* linked to cancer-related cellular processes^82^. Contrary to a predominant mRNA downregulation pattern across cancers^29^, we found that most RBPs genomic alterations in BC are mRNA upregulation (68.7%) and amplification (15.4%) (Fig. 1B). This probably will increase RBPs cellular concentrations leading to dysfunctional post-transcriptional processes.

To determine how many RBPs have been previously studied in BC, we next analyzed the most recent catalog of cancer driver genes: NCG6^30^. Only 14 RBPs have been catalogued as BC driver genes (Fig. 2A). This indicates that RBPs have been poorly investigated in breast carcinogenesis. Thus, to identify new putative BC RBPs, we first explored their genomic alteration profiles associated with tumor progression or suppression (Table 1, Supp. Table 2). As expected, we have found well-known BC progressors and suppressors proteins, such as DAP3^18^, MTDH^19^ or CCAR2^20^, validating our strategy (Table 1). On the contrary, our strategy revealed RBPs that have not been previously associated with tumorigenesis, and yet they are highly altered in BC (e.g. TFB2M, C1ORF131 or DDX19A) (Table 1). Interestingly, the most altered RBP (MRPL13) in our analysis has never been studied in cancer. MRPL13, along with other highly altered RBPs (Table 1), has only shown to interact with ESR2, a tumor suppressor in breast and other cancer types^33^. This observation led us to investigate how many RBPs interact with ESR2; strikingly, we found that 30% of all RBPs interact with this receptor^33^. ESR2 could probably exert its suppressive activity through post-transcriptional mechanisms implicating several RBPs; nevertheless, more research is needed to understand this observation.

Second, to further characterize RBPs associated with BC subtypes and staging, we hence analyzed RBPs genomic alterations (Fig. 2C-D). Interestingly, RBPs genomic alterations gradually increased from Normal to Basal subtype (Fig. 2C), i.e. from low to a high proliferative stage^2^. Concordantly, metastasized tumors (stage IV) showed high frequencies of RBPs genomic alterations compared to non-metastasized samples (stage I to III) (Fig. 2D). Therefore, it seems that RBPs are acting as BC progressors rather than suppressors, which agrees with their genomic alteration profiles (Fig. 1B). This analysis also revealed highly altered RBPs per subtype or staging (Fig. 2C-D; Supp. Table 4-5) that could lead to the discovery of new clinical biomarkers or therapeutic targets. Indeed, SCAMP3 and HEATR6, which have not been studied in BC, presented similar degrees of genomic alterations (Fig. 2D) than well-known metastasis drivers: ARF1^34^ and PARP1^35^. In hepatocellular carcinoma cells, SCAMP3 knockdown has been shown to suppress cell proliferation^83^, while HEATR6 has never been associated with tumorigenesis. Thus, more research is needed to understand their role in BC.

Interaction networks have been shown to be useful for identifying key tumoral proteins^37^. In this regard, by analyzing PPIs between RBPs and well-known BC proteins, we identified SF3B1 and CDC5L at the core of two main networks (Fig. 3). While SF3B1 has been previously implicated in BC^84^, CDC5L, which interacts with 14 BC proteins, has not been studied in this malignancy. However, CDC5L has been related with other cancer types, such as osteosarcoma^85^ and prostate cancer^86^, showing its possible role in breast carcinogenesis.

Next, we exploited the HPA database^14–16^ to identify differentially expressed RBPs in tumor breast tissues. We found 24 upregulated and 62 downregulated RBPs compared with normal tissues. As expected, our approach revealed well-known BC proteins. For instance, KPNA2 which has been known to enhance BC metastasis *ex vivo*^21^, is highly expressed in BC tissues (7 out of 12 tumor samples are classified as high) (Fig. 4B, Supp. Table 6). On the contrary, G3BP2 expression is reduced in tumoral breast tissues (Fig. 4B, Supp. Table 6); accordingly, loss of G3BP2 enhances tumor invasion and metastasis *in vivo*^22^. Interestingly, DARS2 which has never been related with BC is upregulated in our analysis (10 out 12 tumor samples are classified as high) (Fig. 4B, Supp. Table 6) and has been associated with hepatocarcinogenesis progression^87^, showing its putative implication in BC. In addition, SUPT6H protein expression is diminished in breast tumoral tissues (Fig. 4B, Supp. Table 6) and has not been linked to this malignancy. Worthy of note, SUPT6H knockdown is associated with DNA damage via formation of RNA:DNA hybrids (R-loops) in HeLa cells^88^, showing its possible role in breast tumorigenesis.

To identify essential RBPs for tumor survival, we next analyzed *ex vivo* loss-of-function screens: CERES^11^ and DEMETER2^12,13^. *In toto*, we identified 207 essential RBPs for tumor survival. This was expected since RBPs control every aspect of RNA metabolism. However, only 59 were characterized as essential by both computational methods (Fig. 5A; Supp. Table 8-9). Although CERES^11^ and DEMETER2^12,13^ did not tested all human RBPs, therapeutic posttranscriptional BC research could therefore be focused on these 59 RBPs. However, more research is needed to deeply understand their carcinogenic roles. Also, we revealed essential RBPs per BC molecular subtype (Fig. 5B) that could be analyzed to better understand subtype-related posttranscriptional processes.

In extending the scope of our previous analyses, we finally reasoned that integration of all databases examined could narrow down the identification of potential BC RBPs. First, we focused on RBPs having putative tumor progression profiles unraveling 19 RBPs with tumorigenic characteristics according to our analyses (Fig. 5A). As expected, most of them (13 out 19) have been described as BC tumor progressors, controlling different cellular processes: migration, invasion, and metastasis^43–46,49–60^. Interestingly, NSF, SF3A3, PRPF3, and MAGOHB have never been studied in cancer. While on the other hand, PUF60 has been associated with colon and non-small cell lung cancer^47,48^ and PLEC has been shown to promote migration and invasion of neck squamous cell carcinoma^61^. Next, we identified new RBPs having possible tumor suppression roles in BC. As discussed before and depicted in Fig. 1B and Fig. 2 C-D, RBPs seemed to act as progressors rather that suppressors. Although most of the proteins identified have been associated with breast and other cancer types, their suppressive roles remain undetermined ^62–74^. Only 3 have been described as tumor suppressors in BC (CTNND1^23^, LIMA1^24^, AATF^25^), and 4 has been identified as tumor suppressors in other cancer types (MEX3C^75^, UPF1^76^, CNOT1^77^, and TNKS1BP1^78^). Interestingly, 5 have never been studied in cancer: CSTF2T, SRRM1, CPSF7, EIF4G3, and ELOA.

In sum, individual and integrated analysis of the aforementioned databases led us to identify RBPs that have never been studied in BC but with defined tumorigenic functions in other cancer types. Thus, based on their tumorigenic characteristics presented in this study and their roles in other cancer types, we have identified new putative BC RBPs: 6 progressors (MRPL13, SCAMP3, CDC5L, DARS2, PUF60, and PLEC), and 5 suppressors (SUPT6H, MEX3C, UPF1, CNOT1, and TNKS1BP1). However, further research should focus on the mechanisms by which these proteins promote or suppress breast tumorigenesis, holding the promise of new therapeutic pathways along with novel drug development strategies.

## Methods

### Gene sets

A total of 1393 RBPs were extracted form Hentze *et al.*^27^ and checked for new annotations using Ensembl (http://www.ensembl.org)^89,90^. Only one duplicate was found: ENSG00000100101 and ENSG00000273899, both correspond to *NOL12*, leaving a final list of 1392 RBPs. BC genes (n = 171), related to Fig. 1A, were obtained from The Network of Cancer Genes 6.0 (NCG6)^30^. Non-cancer gene list, related to Fig.1A, was constructed as follow: non-cancer genes from Piazza *et al*.^31^, without RBPs and NCG6^30^ cancer genes, were reanalyzed using Piazza’s OncoScore algorithm (http://www.galseq.com/oncoscore.html), giving a final list of 177 non-cancer genes (Supp. Table S1).

### Genomic analysis

Genomic alterations of RBPs, non-cancer and BC genes were analyzed through the cBioPortal (http://www.cbioportal.org/)^91,92^ using the Breast Invasive Carcinoma (TCGA, PanCancer Atlas) database (n = 994 complete samples)^5–10^. To compare the aforementioned gene sets, genomic alterations per protein were corrected by the number of genes or individuals (Fig. 1A, Fig. 2C and D). A Mann–Whitney U test was used to compare genomic alterations between gene sets or clinical characteristics.

### Network construction

Experimental and database interactions (Figure 3 and Table S3) between RBPs (n=1392) and BC proteins (n=171)^30^, having an interaction score of 0.9 (highest confidence), were extracted from STRING database^93^ and visualized using Cytoscape 3.7.1 platform^94^.

### Protein expression analysis

Immunohistological levels of 1212 available RBPs in normal and BC tissues were extracted from Protein Atlas version 18.1 (https://www.proteinatlas.org/)^14–16^. Expression levels of normal tissues were taken from glandular cells, while a consensus level was manually generated for BC tissues (Table S6) based on tissue level frequency. Imunohistological images presented in Figure 4B were taken from https://www.proteinatlas.org/ENSG00000182481-KPNA2/tissue/breast#img (KPNA2 staining in normal tissue), https://www.proteinatlas.org/ENSG00000182481-KPNA2/pathology/tissue/breast+cancer#img (KPNA2 staining in tumoral tissue), https://www.proteinatlas.org/ENSG00000138757-G3BP2/tissue/breast#img (G3BP2 staining in normal tissue), https://www.proteinatlas.org/ENSG00000138757-G3BP2/pathology/breast+cancer#img (G3BP2 staining in tumoral tissue), https://www.proteinatlas.org/ENSG00000109111-SUPT6H/tissue/breast#img (SUPT6H staining in normal tissue), https://www.proteinatlas.org/ENSG00000138757-G3BP2/pathology/breast+cancer#img (SUPT6H staining in tumoral tissue),

### Cancer dependency analysis

RBPs cancer dependency scores from CERES^11^ (1288 available RBPs) and DEMETER2^12,13^ (1290 available RBPs) were obtained from the Dependency Map (DepMap) portal (https://depmap.org/portal/). Molecular subtypes of 82 (DEMETER2^12,13^) and 28 (CERES^11^) BC cell lines were obtained from Smith *et al.*^40^, Dai *et al.*^41^, and Kao *et al.*^42^ (Supp. Table 10)

## Acknowledgements

A.I. acknowledges the Spanish Ministry (MEIC) to EMBL partnership, Severo Ochoa and the CERCA program.

## References

1. Bray, F. et al. Global cancer statistics 2018: GLOBOCAN estimates of incidence and mortality worldwide for 36 cancers in 185 countries. CA. Cancer J. Clin. 68, 394–424 (2018).

2. Harbeck, N. et al. Breast cancer. Nature reviews. Disease primers 5, 66 (2019).

3. Guerrero, S. et al. Analysis of Racial/Ethnic Representation in Select Basic and Applied Cancer Research Studies. Sci. Rep. 8, 13978 (2018).

4. Hiatt, R. A. & Brody, J. G. Environmental Determinants of Breast Cancer. Annu. Rev. Public Health 39, 113–133 (2018).

5. Hoadley, K. A. et al. Cell-of-Origin Patterns Dominate the Molecular Classification of 10,000 Tumors from 33 Types of Cancer. Cell 173, 291-304.e6 (2018).

6. Ellrott, K. et al. Scalable Open Science Approach for Mutation Calling of Tumor Exomes Using Multiple Genomic Pipelines. Cell Syst. (2018). doi:10.1016/j.cels.2018.03.002

7. Taylor, A. M. et al. Genomic and Functional Approaches to Understanding Cancer Aneuploidy. Cancer Cell 33, 676-689.e3 (2018).

8. Gao, Q. et al. Driver Fusions and Their Implications in the Development and Treatment of Human Cancers. Cell Rep. (2018). doi:10.1016/j.celrep.2018.03.050

9. Liu, J. et al. An Integrated TCGA Pan-Cancer Clinical Data Resource to Drive High-Quality Survival Outcome Analytics. Cell (2018). doi:10.1016/j.cell.2018.02.052

10. Sanchez-Vega, F. et al. Oncogenic Signaling Pathways in The Cancer Genome Atlas. Cell 173, 321-337.e10 (2018).

11. Meyers, R. M. et al. Computational correction of copy number effect improves specificity of CRISPR–Cas9 essentiality screens in cancer cells. Nat. Genet. 49, 1779–1784 (2017).

12. Tsherniak, A. et al. Defining a Cancer Dependency Map. Cell 170, 564-576.e16 (2017).

13. McFarland, J. M. et al. Improved estimation of cancer dependencies from large-scale RNAi screens using model-based normalization and data integration. Nat. Commun. 9, 4610 (2018).

14. Uhlén, M. et al. Proteomics. Tissue-based map of the human proteome. Science 347, 1260419 (2015).

15. Thul, P. J. et al. A subcellular map of the human proteome. Science 356, eaal3321 (2017).

16. Uhlen, M. et al. A pathology atlas of the human cancer transcriptome. Science 357, eaan2507 (2017).

17. Abdel-Wahab, O. & Gebauer, F. Editorial overview: Cancer genomics: RNA metabolism and translation in cancer pathogenesis and therapy. Current Opinion in Genetics and Development 48, iv–vi (2018).

18. Wazir, U. et al. Effects of the knockdown of death-associated protein 3 expression on cell adhesion, growth and migration in breast cancer cells. Oncol. Rep. 33, 2575–2582 (2015).

19. Yu, J. et al. MicroRNA-320a inhibits breast cancer metastasis by targeting metadherin. Oncotarget 7, 38612–38625 (2016).

20. Qin, B. et al. DBC1 Functions as a Tumor Suppressor by Regulating p53 Stability. Cell Rep. 10, 1324–1334 (2015).

21. Noetzel, E. et al. Nuclear transport receptor karyopherin-α2 promotes malignant breast cancer phenotypes in vitro. Oncogene 31, 2101–2114 (2012).

22. Wei, S. C. et al. Matrix stiffness drives epithelial-mesenchymal transition and tumour metastasis through a TWIST1-G3BP2 mechanotransduction pathway. Nat. Cell Biol. 17, 678–88 (2015).

23. Schackmann, R. C. J. et al. Loss of p120-catenin induces metastatic progression of breast cancer by inducing anoikis resistance and augmenting growth factor receptor signaling. Cancer Res. 73, 4937–49 (2013).

24. Jiang, W. G. et al. Eplin-alpha expression in human breast cancer, the impact on cellular migration and clinical outcome. Mol. Cancer 7, 71 (2008).

25. Sharma, M. Apoptosis-antagonizing transcription factor (AATF) gene silencing: role in induction of apoptosis and down-regulation of estrogen receptor in breast cancer cells. Biotechnol. Lett. 35, 1561–70 (2013).

26. Lopez-Cortes, A. et al. Prediction of breast cancer proteins using molecular descriptors and artificial neural networks: a focus on cancer immunotherapy proteins, metastasis driver proteins, and RNA-binding proteins. bioRxiv Bioinforma. (2019). doi:10.1101/840108

27. Hentze, M. W., Castello, A., Schwarzl, T. & Preiss, T. A brave new world of RNA-binding proteins. Nat. Rev. Mol. Cell Biol. 19, 327–341 (2018).

28. Wang, K. et al. Integrated Bioinformatics Analysis the Function of RNA Binding Proteins (RBPs) and Their Prognostic Value in Breast Cancer. Front. Pharmacol. 10, 140 (2019).

29. Wang, Z. L. et al. Comprehensive Genomic Characterization of RNA-Binding Proteins across Human Cancers. Cell Rep. 22, 286–298 (2018).

30. Repana, D. et al. The Network of Cancer Genes (NCG): a comprehensive catalogue of known and candidate cancer genes from cancer sequencing screens. Genome Biol. 20, 1 (2019).

31. Piazza, R. et al. OncoScore: a novel, Internet-based tool to assess the oncogenic potential of genes. Sci. Rep. 7, 46290 (2017).

32. Wurth, L. et al. UNR/CSDE1 Drives a Post-transcriptional Program to Promote Melanoma Invasion and Metastasis. Cancer Cell 30, 694–707 (2016).

33. Giurato, G. et al. Quantitative mapping of RNA-mediated nuclear estrogen receptor β interactome in human breast cancer cells. Sci. Data 5, 180031 (2018).

34. Xie, X. et al. Suppression of breast cancer metastasis through the inactivation of ADP-ribosylation factor 1. Oncotarget 7, 58111–58120 (2016).

35. Zimmer, A. S., Gillard, M., Lipkowitz, S. & Lee, J.-M. Update on PARP Inhibitors in Breast Cancer. Curr. Treat. Options Oncol. 19, 21 (2018).

36. Rodríguez, M. I. et al. PARP-1 Regulates Metastatic Melanoma through Modulation of Vimentin-induced Malignant Transformation. PLoS Genet. 9, e1003531 (2013).

37. Wurth, L. et al. UNR/CSDE1 Drives a Post-transcriptional Program to Promote Melanoma Invasion and Metastasis. Cancer Cell 30, 694–707 (2016).

38. Szklarczyk, D. et al. STRING v11: Protein-protein association networks with increased coverage, supporting functional discovery in genome-wide experimental datasets. Nucleic Acids Res. 47, D607–D613 (2019).

39. Reimand, J., Kull, M., Peterson, H., Hansen, J. & Vilo, J. g:Profiler--a web-based toolset for functional profiling of gene lists from large-scale experiments. Nucleic Acids Res. 35, W193–200 (2007).

40. Smith, S. E. et al. Molecular characterization of breast cancer cell lines through multiple omic approaches. Breast Cancer Res. 19, 65 (2017).

41. Dai, X., Cheng, H., Bai, Z. & Li, J. Breast Cancer Cell Line Classification and Its Relevance with Breast Tumor Subtyping. J. Cancer 8, 3131–3141 (2017).

42. Kao, J. et al. Molecular Profiling of Breast Cancer Cell Lines Defines Relevant Tumor Models and Provides a Resource for Cancer Gene Discovery. PLoS One 4, e6146 (2009).

43. Jian, J., Yang, Q. & Huang, X. Src Regulates Tyr ^20^ Phosphorylation of Transferrin Receptor-1 and Potentiates Breast Cancer Cell Survival. J. Biol. Chem. 286, 35708–35715 (2011).

44. Singh, M. et al. Differential Expression of Transferrin Receptor (TfR) in a Spectrum of Normal to Malignant Breast Tissues. Appl. Immunohistochem. Mol. Morphol. 19, 417–423 (2011).

45. Habashy, H. O. et al. Transferrin receptor (CD71) is a marker of poor prognosis in breast cancer and can predict response to tamoxifen. Breast Cancer Res. Treat. 119, 283–293 (2010).

46. Sheng, C. et al. Suppression of Kpnβ1 expression inhibits human breast cancer cell proliferation by abrogating nuclear transport of Her2. Oncol. Rep. 39, 554–564 (2017).

47. Kobayashi, S. et al. Anti-FIRs (PUF60) auto-antibodies are detected in the sera of early-stage colon cancer patients. Oncotarget 7, 82493–82503 (2016).

48. Müller, B. et al. Concomitant expression of far upstream element (*FUSE*) binding protein (*FBP*) interacting repressor (FIR) and its splice variants induce migration and invasion of non-small cell lung cancer (NSCLC) cells. J. Pathol. 237, 390–401 (2015).

49. Wazir, U. et al. Effects of the knockdown of death-associated protein 3 expression on cell adhesion, growth and migration in breast cancer cells. Oncol. Rep. 33, 2575–2582 (2015).

50. Ray Chaudhuri, A. & Nussenzweig, A. The multifaceted roles of PARP1 in DNA repair and chromatin remodelling. Nat. Rev. Mol. Cell Biol. 18, 610–621 (2017).

51. Wang, Y. et al. Tricho-rhino-phalangeal syndrome 1 protein functions as a scaffold required for ubiquitin-specific protease 4-directed histone deacetylase 2 deubiquitination and tumor growth. Breast Cancer Res. 20, 83 (2018).

52. Montanaro, L. et al. Relationship between dyskerin expression and telomerase activity in human breast cancer. Cell. Oncol. 30, 483–90 (2008).

53. Jiang, B.-H. et al. Poly(ADP-Ribose) Polymerase 1: Cellular Pluripotency, Reprogramming, and Tumorogenesis. Int. J. Mol. Sci. 16, 15531–15545 (2015).

54. Dedes, K. J. et al. Cortactin gene amplification and expression in breast cancer: a chromogenic in situ hybridisation and immunohistochemical study. Breast Cancer Res. Treat. 124, 653–666 (2010).

55. Wang, N., Ma, Q., Wang, Y., Ma, G. & Zhai, H. CacyBP/SIP Expression is Involved in the Clinical Progression of Breast Cancer. World J. Surg. 34, 2545–2552 (2010).

56. Ridgway, L. D., Wetzel, M. D., Ngo, J. A., Erdreich-Epstein, A. & Marchetti, D. Heparanase-Induced GEF-H1 Signaling Regulates the Cytoskeletal Dynamics of Brain Metastatic Breast Cancer Cells. Mol. Cancer Res. 10, 689–702 (2012).

57. Sevinsky, C. J. et al. NDRG1 regulates neutral lipid metabolism in breast cancer cells. Breast Cancer Res. 20, 55 (2018).

58. Ma, A. et al. USP1 inhibition destabilizes KPNA2 and suppresses breast cancer metastasis. Oncogene 38, 2405–2419 (2019).

59. Zhu, S., Sachdeva, M., Wu, F., Lu, Z. & Mo, Y.-Y. Ubc9 promotes breast cell invasion and metastasis in a sumoylation-independent manner. Oncogene 29, 1763–1772 (2010).

60. Moela, P., Choene, M. M. S. & Motadi, L. R. Silencing RBBP6 (Retinoblastoma Binding Protein 6) sensitises breast cancer cells MCF7 to staurosporine and camptothecin-induced cell death. Immunobiology 219, 593–601 (2014).

61. Katada, K. et al. Plectin promotes migration and invasion of cancer cells and is a novel prognostic marker for head and neck squamous cell carcinoma. J. Proteomics 75, 1803–1815 (2012).

62. Fei, T. et al. Genome-wide CRISPR screen identifies HNRNPL as a prostate cancer dependency regulating RNA splicing. Proc. Natl. Acad. Sci. U. S. A. 114, E5207–E5215 (2017).

63. Panagopoulos, I. et al. Fusion of the SEC31L1 and ALK genes in an inflammatory myofibroblastic tumor. Int. J. cancer 118, 1181–6 (2006).

64. Hinrichsen, I. et al. Reduced migration of MLH1 deficient colon cancer cells depends on SPTAN1. Mol. Cancer 13, 11 (2014).

65. Hubaux, R. et al. Microtubule affinity-regulating kinase 2 is associated with DNA damage response and cisplatin resistance in non-small cell lung cancer. Int. J. cancer 137, 2072–82 (2015).

66. Kim, M. S. et al. Somatic mutations and losses of expression of microRNA regulation-related genes AGO2 and TNRC6A in gastric and colorectal cancers. J. Pathol. 221, 139–46 (2010).

67. Wang, Y. et al. The mismatch repair gene hPMS1 (human postmeiotic segregation1) is down regulated in oral squamous cell carcinoma. Gene 524, 28–34 (2013).

68. Degot, S. et al. Association of the breast cancer protein MLN51 with the exon junction complex via its speckle localizer and RNA binding module. J. Biol. Chem. 279, 33702–15 (2004).

69. Tyson-Capper, A. & Gautrey, H. Regulation of Mcl-1 alternative splicing by hnRNP F, H1 and K in breast cancer cells. RNA Biol. 15, 1448–1457 (2018).

70. Cruz-Ramos, E., Macías-Silva, M., Sandoval-Hernández, A. & Tecalco-Cruz, A. C. Non-muscle myosin IIA is post-translationally modified by interferon-stimulated gene 15 in breast cancer cells. Int. J. Biochem. Cell Biol. 107, 14–26 (2019).

71. Singel, S. M. et al. A targeted RNAi screen of the breast cancer genome identifies KIF14 and TLN1 as genes that modulate docetaxel chemosensitivity in triple-negative breast cancer. Clin. Cancer Res. 19, 2061–70 (2013).

72. Iyer, R. S. et al. The RNA helicase/transcriptional co-regulator, p68 (DDX5), stimulates expression of oncogenic protein kinase, Polo-like kinase-1 (PLK1), and is associated with elevated PLK1 levels in human breast cancers. Cell Cycle 13, 1413–1423 (2014).

73. Schatz, N., Brändlein, S., Rückl, K., Hensel, F. & Vollmers, H. P. Diagnostic and therapeutic potential of a human antibody cloned from a cancer patient that binds to a tumor-specific variant of transcription factor TAF15. Cancer Res. 70, 398–408 (2010).

74. Badura, M., Braunstein, S., Zavadil, J. & Schneider, R. J. DNA damage and eIF4G1 in breast cancer cells reprogram translation for survival and DNA repair mRNAs. Proc. Natl. Acad. Sci. U. S. A. 109, 18767–72 (2012).

75. Burrell, R. A. et al. Replication stress links structural and numerical cancer chromosomal instability. Nature 494, 492–496 (2013).

76. Li, L. et al. The Human RNA Surveillance Factor UPF1 Modulates Gastric Cancer Progression by Targeting Long Non-Coding RNA MALAT1. Cell. Physiol. Biochem. 42, 2194–2206 (2017).

77. Vicente, C. et al. The CCR4-NOT complex is a tumor suppressor in Drosophila melanogaster eye cancer models. J. Hematol. Oncol. 11, 108 (2018).

78. Ohishi, T. et al. Tankyrase-Binding Protein TNKS1BP1 Regulates Actin Cytoskeleton Rearrangement and Cancer Cell Invasion. Cancer Res. 77, 2328–2338 (2017).

79. López-Cortés, A. et al. OncoOmics approaches to reveal essential genes in breast cancer: a panoramic view from pathogenesis to precision medicine. bioRxiv 638866 (2019). doi:10.1101/638866

80. López-Cortés, A. et al. Gene prioritization, communality analysis, networking and metabolic integrated pathway to better understand breast cancer pathogenesis. Sci. Rep. 8, 16679 (2018).

81. López-Cortés, A. et al. Prediction of druggable proteins using machine learning and functional enrichment analysis: a focus on cancer-related proteins and RNA-binding proteins. bioRxiv 825513 (2019). doi:10.1101/825513

82. Moore, S., Järvelin, A. I., Davis, I., Bond, G. L. & Castello, A. Expanding horizons: new roles for non-canonical RNA-binding proteins in cancer. Current Opinion in Genetics and Development 48, 112–120 (2018).

83. Zhang, X. et al. Overexpression of SCAMP3 is an indicator of poor prognosis in hepatocellular carcinoma. Oncotarget 8, 109247–109257 (2017).

84. Maguire, S. L. et al. SF3B1 mutations constitute a novel therapeutic target in breast cancer. J. Pathol. 235, 571–580 (2015).

85. Lu, X.-Y. et al. Cell Cycle Regulator Gene CDC5L, a Potential Target for 6p12-p21 Amplicon in Osteosarcoma. Mol. Cancer Res. 6, 937–946 (2008).

86. Li, X. et al. Oncogenic Properties of NEAT1 in Prostate Cancer Cells Depend on the CDC5L–AGRN Transcriptional Regulation Circuit. Cancer Res. 78, 4138–4149 (2018).

87. Qin, X. et al. Upregulation of DARS2 by HBV promotes hepatocarcinogenesis through the miR-30e-5p/MAPK/NFAT5 pathway. J. Exp. Clin. Cancer Res. 36, 148 (2017).

88. Nojima, T. et al. Deregulated Expression of Mammalian lncRNA through Loss of SPT6 Induces R-Loop Formation, Replication Stress, and Cellular Senescence. Mol. Cell 72, 970-984.e7 (2018).

89. Hunt, S. E. et al. Ensembl variation resources. Database 2018, (2018).

90. Zerbino, D. R. et al. Ensembl 2018. Nucleic Acids Res. 46, D754–D761 (2018).

91. Cerami, E. et al. The cBio cancer genomics portal: an open platform for exploring multidimensional cancer genomics data. Cancer Discov. (2012). doi:10.1158/2159-8290.CD-12-0095

92. Gao, J. et al. Integrative analysis of complex cancer genomics and clinical profiles using the cBioPortal. Sci. Signal. (2013). doi:10.1126/scisignal.2004088

93. Szklarczyk, D. et al. STRING v10: protein-protein interaction networks, integrated over the tree of life. Nucleic Acids Res. 43, D447–52 (2015).

94. Shannon, P. et al. Cytoscape: a software environment for integrated models of biomolecular interaction networks. Genome Res. 13, 2498–504 (2003).

95. Chin, S. F. et al. High-resolution aCGH and expression profiling identifies a novel genomic subtype of ER negative breast cancer. Genome Biol. 8, R215 (2007).

96. Cao, J. et al. The overexpression and prognostic role of DCAF13 in hepatocellular carcinoma. Tumor Biol. 39, 101042831770575 (2017).

97. Li, Y. et al. Amplification of LAPTM4B and YWHAZ contributes to chemotherapy resistance and recurrence of breast cancer. Nat. Med. 16, 214–218 (2010).

98. Chang, C.-C. et al. Upregulation of lactate dehydrogenase a by 14-3-3ζ leads to increased glycolysis critical for breast cancer initiation and progression. Oncotarget 7, 35270–83 (2016).

99. Jacques, C. et al. Death-associated protein 3 is overexpressed in human thyroid oncocytic tumours. Br. J. Cancer 101, 132–138 (2009).

100. Drosos, Y. et al. NUCKS overexpression in breast cancer. Cancer Cell Int. 9, 19 (2009).

101. Shi, C. et al. NUCKS nuclear elevated expression indicates progression and prognosis of ovarian cancer. Tumor Biol. 39, 101042831771463 (2017).

102. Li, J. et al. Knockdown of metadherin inhibits cell proliferation and migration in colorectal cancer. Oncol. Rep. 40, 2215–2223 (2018).

103. Best, S. A., Nwaobasi, A. N., Schmults, C. D. & Ramsey, M. R. CCAR2 Is Required for Proliferation and Tumor Maintenance in Human Squamous Cell Carcinoma. J. Invest. Dermatol. 137, 506–512 (2017).

104. Jen, J. et al. Oncoprotein ZNF322A transcriptionally deregulates alpha-adducin, cyclin D1 and p53 to promote tumor growth and metastasis in lung cancer. Oncogene 35, 2357–2369 (2016).

105. Gala, K. et al. KMT2C mediates the estrogen dependence of breast cancer through regulation of ERα enhancer function. Oncogene 37, 4692–4710 (2018).

106. Xia, M. et al. Downregulation of MLL3 in esophageal squamous cell carcinoma is required for the growth and metastasis of cancer cells. Tumor Biol. 36, 605–613 (2015).

107. Kim, S. et al. Blocking Myristoylation of Src Inhibits Its Kinase Activity and Suppresses Prostate Cancer Progression. Cancer Res. 77, 6950–6962 (2017).

